# Substrate promiscuity of xenobiotic-transforming hydrolases from stream biofilms impacted by treated wastewater

**DOI:** 10.1101/2023.09.27.559296

**Authors:** Yaochun Yu, Niklas Ferenc Trottmann, Milo R. Schärer, Kathrin Fenner, Serina L. Robinson

## Abstract

Organic contaminants enter aquatic ecosystems from various sources, including wastewater treatment plant effluent. Freshwater biofilms play a major role in the removal of organic contaminants from receiving water bodies, but knowledge of the molecular mechanisms driving contaminant biotransformations in complex stream biofilm (periphyton) communities remains limited. Previously, we demonstrated that biofilms in experimental flume systems grown at higher ratios of treated wastewater (WW) to stream water displayed an increased biotransformation potential for a number of organic contaminants. We identified a positive correlation between WW percentage and biofilm biotransformation rates for the widely-used insect repellent, *N,N*-diethyl-meta-toluamide (DEET). Here, we conducted deep shotgun sequencing of flume biofilms and identified a positive correlation between WW percentage and metagenomic read abundances of DEET hydrolase (DH) homologs. To test the causality of this association, we constructed a targeted metagenomic library of DH homologs from flume biofilms. We screened our complete metagenomic library for activity with four different substrates and a subset thereof with 183 WW-related organic compounds. The majority of active hydrolases in our library preferred aliphatic and aromatic ester substrates while, remarkably, only a single reference enzyme was capable of DEET hydrolysis. Of the 626 total enzyme-substrate combinations tested, approximately 5% were active enzyme-substrate pairs. Metagenomic DH family homologs revealed a broad substrate promiscuity spanning 22 different compounds when summed across all enzymes tested. We biochemically characterized the most promiscuous and active enzymes identified based on metagenomic analysis from uncultivated *Rhodospirillaceae* and *Planctomycetaceae*. In addition to characterizing new DH family enzymes, we exemplified a framework for linking metagenome-guided hypothesis generation with experimental validation. Overall, this study expands the scope of known enzymatic contaminant biotransformations for metagenomic hydrolases from WW-receiving stream biofilm communities.

**Graphical abstract:** 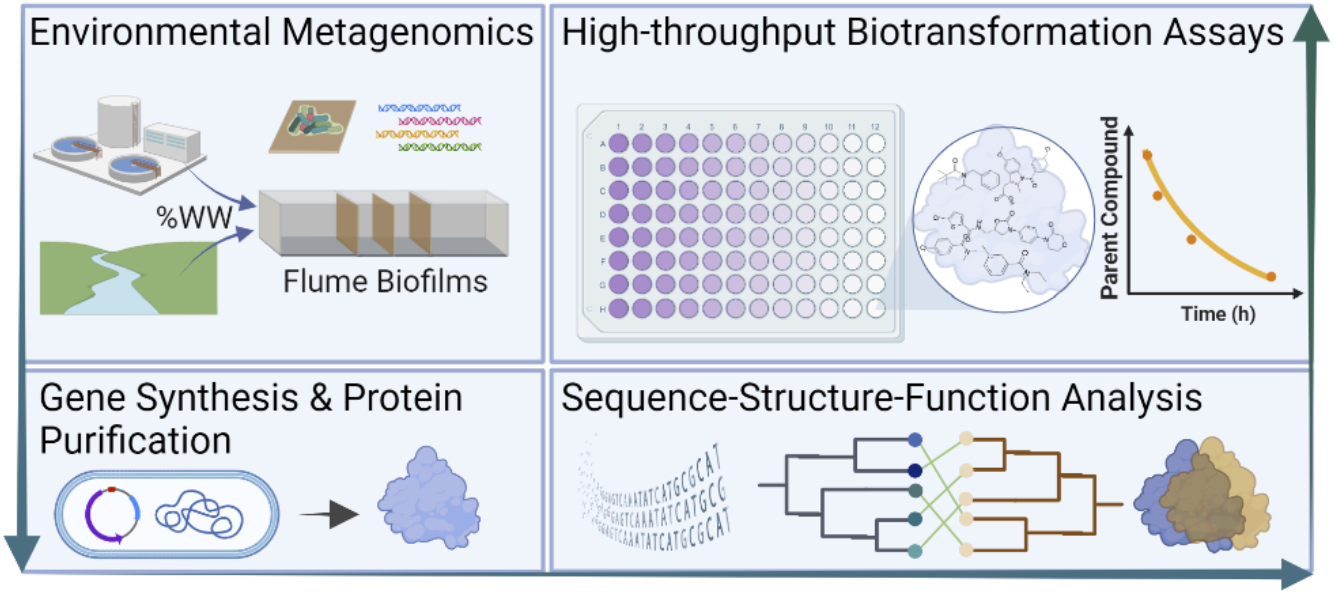

**Highlights:**

- Metagenomic DEET hydrolase abundances higher in biofilms grown in treated wastewater.
- Eleven out of 64 metagenomic hydrolases tested exhibited hydrolysis activity.
- Related enzymes in a single family of DEET hydrolases biotransform 20+ contaminants.
- Reference DEET hydrolase shows a substrate preference for benzamide moieties.
- ‘True’ DEET hydrolases are in low abundance even in biofilms that degrade DEET.

## 1. Introduction

Trace organic contaminants (TrOCs) released due to anthropogenic activities such as industrial production and agricultural practices are exerting increasing pressures on ecosystems and human health (Caliman and Gavrilescu, 2009; Daughton and Ternes, 1999; Evich et al., 2022; Fenner et al., 2013; Halden, 2011; Schwarzenbach et al., 2006). Many TrOCs and their transformation products exert toxicity through bioaccumulation, mixture effects, and specific toxic interactions (Altenburger et al., 2018; Carles et al., 2021; Jin et al., 2023). In aquatic systems, microbes are the dominant players in the biotransformation and removal of various TrOCs through metabolic and co-metabolic processes (Che et al., 2021; Fenner et al., 2013; Han et al., 2023, 2019; Xing et al., 2018; Yu et al., 2018). However, apart from a few specific pathways, the diversity of enzymes involved in microbial biotransformation processes remain largely unknown. Understanding the scope and diversity of microbial enzyme-substrate interactions within environmental microbiomes is critical to improve prediction of biotransformations and promote contaminant removal (Fenner et al., 2021).

Freshwater biofilms are ideal systems for investigating TrOC biotransformation processes due to their complex community composition and critical ecological functions in aquatic environments (Battin et al., 2016; Desiante et al., 2021). Most studies on biofilms have been conducted using amplicon sequencing to profile taxonomic community shifts that correlate with TrOC biotransformations in treated wastewater (Carles et al., 2022, 2021; Desiante et al., 2022; Mansfeldt et al., 2020; Tien et al., 2013). Metagenomics and metatranscriptomics analyses have recently deepened the resolution of community analysis to identify functional shifts in TrOC-exposed engineered wetland biomats (Vega et al., 2023, 2022). However, the number of studies conducted on the molecular mechanisms of TrOC biotransformations in freshwater biofilms or biomats is small relative to other systems such as enriched soil consortia (Vasileiadis et al., 2022) or activated sludge (Rich and Helbling, 2023). Activated sludge studies have used metagenomics and metatranscriptomics to correlate microbial taxa, functional genes, and pollutant biotransformations (Achermann et al., 2020; Athanasakoglou and Fenner, 2022; Kennes-Veiga et al., 2022; Stadler et al., 2018), but also have remained limited in uncovering experimentally-validated enzymes involved in TrOC biotransformations. Many enzymes involved in the biotransformation of contaminants remain a ‘black box.’

In our previous study, stream biofilms cultivated at different ratios of stream water and either ultrafiltered or non-ultrafiltered treated wastewater (WW) exhibited varying TrOC biotransformation capacities (Desiante et al., 2022). Among 75 TrOCs tested, seven compounds (*N,N*-diethyl-meta-toluamide (DEET), acesulfame K, caffeine, cyclamate, levetiracetam, saccharin, and valsartan, Figure S1) showed clear evidence for increased biotransformation rates in biofilms grown at higher WW percentages, as compared to controls grown in stream water or in ultrafiltered WW. This phenomenon, previously defined as the *downstream effect* (Desiante et al., 2022), strongly suggested that microorganisms present in the WW might affect the biotransformation capacity of biofilms downstream of wastewater treatment plants. Notably, all compounds that exhibited the *downstream effect* contained hydrolyzable moieties, primarily sulfonamide- and amide-functional groups (Figure S1). This motivated us to initiate a large-scale investigation into the potential roles of environmental hydrolases from wastewater effluent-impacted biofilms in degrading TrOCs.

Here we employed a metagenome mining framework to investigate contaminant biotransformations and enzyme-substrate specificity in complex stream biofilm communities. We observed that increasing biotransformation rates of DEET (an insect repellant and one of the seven *downstream effect* compounds) corresponded with higher metagenomic abundances of DEET hydrolase (DH) homologs in stream biofilms with increasing proportions of treated wastewater (WW). Because of the similarity in hydrolyzable moieties amongst all *downstream effect* compounds, we hypothesized that diverse metagenomic DH family homologs detected in WW-impacted biofilms have an expanded substrate specificity not only for DEET, but also for other WW-relevant contaminants. Guided by protein structural modeling from metagenomic sequences, we designed, constructed, and tested a targeted metagenomic library of 64 DH homologs. This work maps the sequence-structure-substrate relationships of DH family enzymes encoded by ten different bacterial phyla from complex stream biofilm communities. The results not only expand our fundamental understanding of how enzyme diversity relates to biotransformation processes, but also have implications for the identification of novel and functional TrOC-degrading enzymes from environmental microbiomes.

## 2. Materials and Methods

### 2.1. Biofilm sampling

The biofilm samples which in this study were used for metagenomic sequencing and hydrolase screening were previously analyzed by 16S and 18S rRNA gene amplicon sequencing (Carles et al., 2021) and organic contaminant biotransformation experiments (Desiante et al., 2022). Briefly, the experimental flume systems (Burdon et al., 2020) for biofilm growth consisted of two independent experiments described by (Desiante et al., 2022), with 16 (for experiment 1) and 20 water channels (for experiment 2) fed with different proportions of the following sources: stream water mixed in proportions with either 10%, 30%, or 80% WW (for experiment 1) or 30% or 80% ultrafiltered (pore size 0.4 µm) WW in experiment 2. Biofilms were grown on glass slides placed in these flow-through channels for four weeks. Biofilms from different glass slides within the same channel were scraped independently to form biological replicates (n=4 for each treatment) and suspended in 400 mL of artificial river water (Carles et al., 2022, 2021; Desiante et al., 2022).

### 2.2. DEET hydrolase metagenomic library construction

Metagenomic sequencing and standard pre-processing and assembly pipelines used are described by Attrah et al. (2023). Protein FASTA files from the 75 metagenomes (Table S1) were queried for hits to the sole characterized reference DEET hydrolase (A3R4E0_PSEPU). Filtering steps for metagenomic library selection are described in the SI. The 64 selected metagenomic DH homolog sequences were codon-optimized for *E. coli* expression using the build-optimization software, BOOST (Oberortner et al., 2017), and cloned by the U.S. Department of Energy Joint Genome Institute with C-terminal 6x-His tags, a tobacco etch virus cleavage site, and additional flexible linker into the first multiple cloning site of pCDF-Duet vectors retaining the NdeI and HindIII restriction sites. Sequence-verified constructs were transformed into T7 Express Competent *E. coli* cells (NEB C2566I).

### 2.3. Sequence and structure bioinformatic analysis

Read mapping and coverage calculations for the *P. putida* reference DH and full-length DH homologs from the above-described metagenomic library were computed as described in the SI. DH phylogenetic analysis is further described in the SI. Full-length sequences of the 64 metagenomic DH proteins and reference enzymes were used as inputs for structural modeling using AlphaFold 2.0 ColabFold v1.3.0 (Mirdita et al., 2022) with default parameters. Structural alignment methods are described in the SI.

### 2.4. Colorimetric activity screening of *P. putida* DH and DH homologs

Activity screening for the 64 metagenomic DH homologs and the *P. putida* reference sequence was performed using a previously established colorimetric screening protocol for hydrolase activity (Robinson et al., 2020; Smith et al., 2020) and described in the SI.

### 2.5. HPLC analysis of *P. putida* DH and metagenomic DH homologs hydrolysis activity

The enzyme activity of *P. putida* DH and DH homologs with DEET and methylbenzoate (MB) were tested in crude extracts containing soluble proteins. DEET and MB (final conc. 250 μM) were added to individual vials containing crude extract separately. Subsequently, reaction progress was monitored by sampling and quenching the reaction with an equal volume of acetonitrile at defined time intervals at 0, 1, 3, 5 and 24h and measured by HPLC (Summit HPLC system, Dionex). Detailed information regarding the HPLC method is included in the SI.

### 2.6. Purification of *P. putida* DH and metagenomic DH homologs

Induction and expression of DH homologs, cell lysis, and harvesting of cell pellets were performed as described in the SI with 400 mL Terrific broth culture media (12 g/L tryptone, 24 g/L yeast extract, 0.4% (v/v) glycerol) supplemented with 10% (vol/vol) phosphate buffer (0.17 M KH_2_PO_4_ and 0.72 M K_2_HPO_4_). Protein purification was performed using benchtop purification for initial tests and FPLC (ÄKTA Pure 25, Cytiva) for data and figures reported in the manuscript. Fractions obtained from both benchtop and FPLC purifications were subjected to analysis by SDS-PAGE (Figure S2 – S4). The detailed purification methods can be found in the SI.

### 2.7. TrOCs selection

A total of 183 TrOCs were chosen for biotransformation experiments with purified *P. putida* DH and the highly active DH homolog p055 selected as described in the Results & Discussion. These TrOCs are commonly used pesticides, pharmaceuticals, and artificial sweeteners and were selected to equally cover a number of relevant enzymatic transformation reactions. Stock solutions of TrOCs were individually prepared in DMSO at a concentration of 10 mM and stored at −20°C until use. Detailed information including the suppliers of each compound can be found in Table S4 and S5

### 2.8. High-throughput biotransformation experiments of TrOCs in 96-well plates

Purified *P. putida* DH and the metagenomic DH homolog p055 were tested for their activity with 14 sub-mixtures composed of the 183 TrOCs instead of testing individual compounds. To avoid potential cumulative toxicities and competitive inhibition, we first conducted a hierarchical clustering of all compounds based on their chemical fingerprints and reaction rules triggered by enviPath (Wicker et al., 2016). Sub-mixtures were then designed to contain compounds with different annotated reaction rules. Each of the 14 sub-mixtures (200 μM for each TrOC) was prepared via appropriate dilution of the stock solutions in EtOH (Table S4 and S5) and stored at −20°C for later biotransformation experiments. The capability of TrOCs biotransformation by different enzymes was investigated in a clear-bottomed 96-well plate (Greiner). Individual TrOC sub-mixtures were first added to an empty 96-well plate and the organic solvent EtOH was completely evaporated before 245 μL 10 mM Tris-HCl buffer (pH=8) was added to each well. The plate was shaken at 50 rpm for 15 min to redissolve the TrOCs. Individually purified enzymes (final concentration of ∼190 nM) were then added to a well plate and mixed thoroughly. Samples for LC-HRMS/MS analysis were collected at 0, 1, 6, and 24h. At each sampling event, a 20 μL sample was taken and mixed with the same amount of pure methanol to completely denature the enzyme, 160 μL buffer was then added to the sample mixture for an approximately 10 times dilution of the original sample. After centrifugation (13,000 × g, 4°C for 15 min), the supernatant was collected and stored at 4°C for LC-HRMS/MS analysis. An enzyme-free abiotic fresh buffer control and trichloroacetic acid (TCA) treated inactivated enzyme control were set up in the same way as the biological experiment described above. All experiments were conducted in duplicate. Samples were submitted to LC-HRMS/MS analysis within two days after collection. Matrix match calibration standards were used for quantification. Quantification was based on concentrations calculated from peak area integration using TraceFinder 5.1 (Thermo Fisher Scientific).

### 2.9. UHPLC-HRMS/MS analysis

The parent compounds and transformation products (TPs) were analyzed by UHPLC-HRMS/MS (Q Exactive, Thermo Fisher Scientific) using a method as previously described but with slight modifications (Yu et al., 2022). Detailed methods for the LC method can be found in the SI. Xcalibur 4.0 (Thermo Fisher Scientific) was used for data acquisition and analysis. Suspect screening for DEET transformation products was conducted using TraceFinder 5.1. ChemDraw Professional 20.0 and MarvinSketch (v19.20.0, ChemAxon, http://www.chemaxon.com) were used for drawing, displaying, and characterizing chemical structures. The minimum incubation time for different control groups at which the maximum removal percentage of individual compounds was achieved was calculated based on Equation 1. The results were included in Table S4 (for DEET hydrolase) and Table S5 (for p055). Significant biological removal was defined as following criteria: (1) > 20% removal in enzymatic control, and (2) at least 10% more removal compared to both abiotic buffer control and inactivated enzyme control.

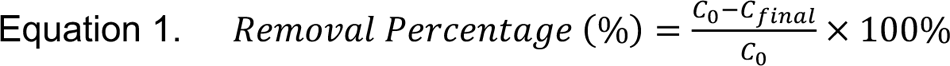

Where C_0_ is the initial concentration measured by LC-HRMS/MS and C_final_ is the final concentration after biotransformation experiments.

## 3. Results and Discussion

### 3.1. Metagenomic sequencing and analysis of WW-impacted biofilm communities

We initiated this study to identify enzymes involved in the biotransformation of TrOCs exhibiting the *downstream effect* in WW-impacted stream biofilms. Our previous study employed 16S rRNA gene analysis to identify differentially-abundant amplicon sequence variants correlating with these TrOC biotransformations (Desiante et al., 2022). However, since 16S rRNA gene sequencing doesn’t capture functional genes, here we used deeply-sequenced metagenomes (up to 200 million reads per sample) from the same biofilms (Attrah et al., 2023) and searched for genes linked to the biodegradation of *downstream effect* TrOCs. Among the proposed biotransformation pathways for the seven TrOCs showing a *downstream effect* (Desiante et al., 2022), only one compound (DEET) could be linked to an experimentally validated hydrolase sequence at the time of our analysis (Rivera-Cancel et al., 2007). The sequence of the protein involved in hydrolysis of the *downstream effect* compound cyclamate remained unknown although its partial purification was reported (Nimura et al., 1974). Several other pathways for *downstream effect* compounds were predicted (Desiante et al., 2022), but only one additional validated hydrolase for the *downstream effect* compound, acesulfame K, was reported during the course of this analysis (Bonatelli et al., 2023). Using the reported acesulfame K hydrolase gene as a query (Bonatelli et al., 2023), we detected no significant BLAST hits in our assembled metagenomes. However, in the same metagenomes we detected more than 3,500 DEET hydrolase (DH) homologs. This diversity and abundance motivated us to investigate experimentally the link between DH homologs and the DEET biotransformation phenotype observed in the biofilms.

Next, we calculated the abundances of DH homologs in assembled metagenomes of biofilms grown in different WW percentages. We used the *Psuedomonas putida* DH as a query since this is the only characterized enzyme known to have DH activity. We quantified sequence counts in the biofilms grown in 0%, 10%, 30%, and 80% treated WW mixed with stream water for two independent flume experiments (Figure 1A). Notably, in two separate, independent flume experiments, we observed a positive correlation between the metagenomic DH abundances in biofilms and the percentage of unfiltered WW the biofilms were grown in (Figure 1A, R^2^ = 0.77 for Exp. 2, R^2^ = 0.42 for Exp. 1). We also calculated the coverage of metagenomic reads mapping to the *P. putida* DH reference in the stream, filtered WW, and unfiltered WW samples and identified significantly more reads mapping in WW relative to stream water (Figure S5, *p*-value < 2e-16).

**Figure 1:**
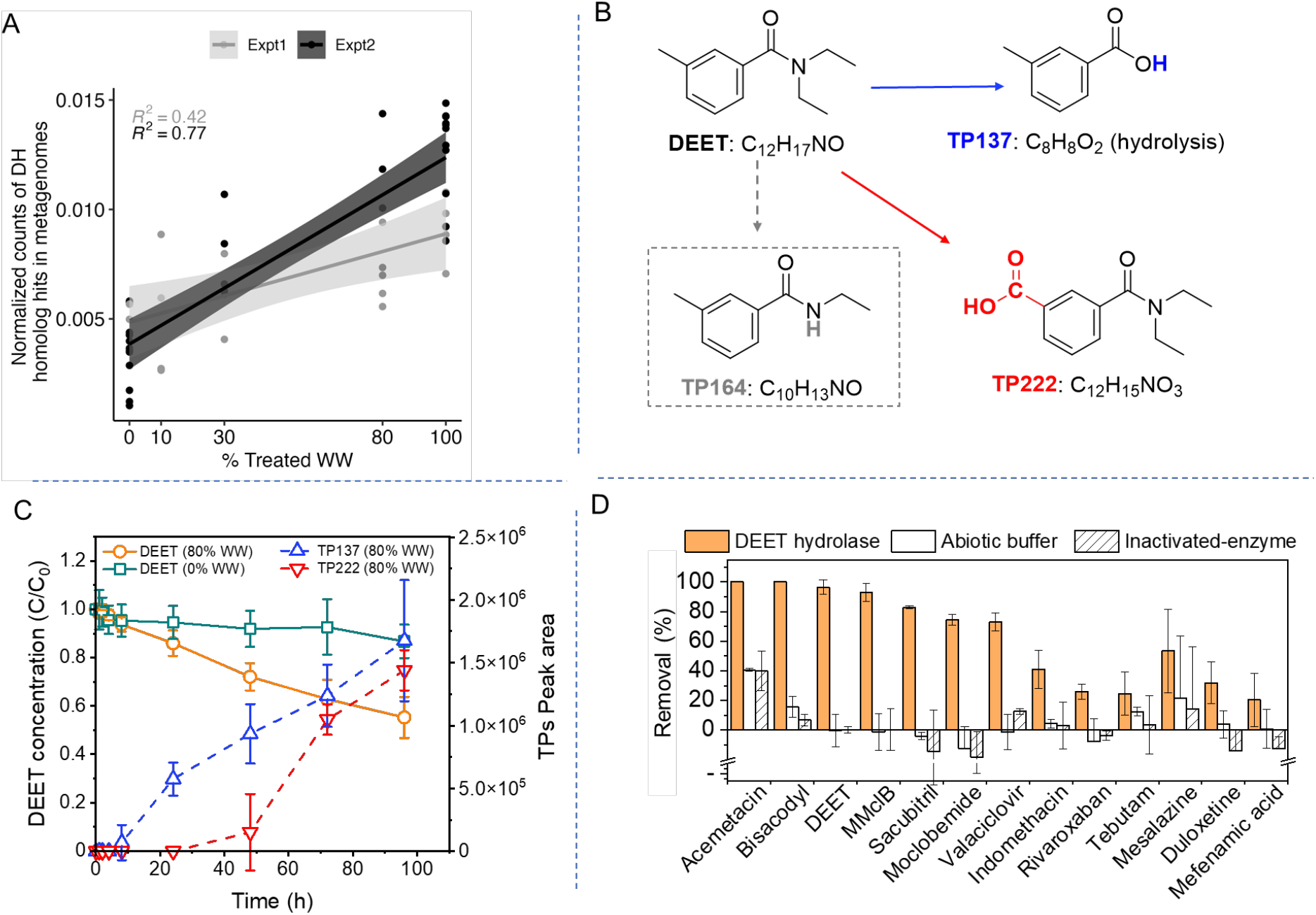
A) Normalized counts of hits in biofilm metagenomes bearing sequence similarity to the *P. putida* DH recovered from assembled metagenomic sequences fitted with a linear regression model (95% confidence interval) across two independent experiments: experiment 1 (light gray) and experiment 2 (dark gray). B) Documented DEET biotransformation pathways and transformation products (TPs) detected in this study. The product TP164 in the dashed-gray box was reported previously (Helbling et al., 2010b) but not detected in this study. C) DEET degradation (solid line) and detected transformation products (TPs) (dashed line) in 80% WW and 0% WW (no TP was detected in 0% WW samples). D) Purified *P. putida* DH substrate specificity for active enzyme (in orange) relative to an inactivated enzyme (hashed lines) and abiotic buffer control (white). Detailed selection criteria and complete biotransformation data are available in the Methods and Table S4. Compounds structures are displayed in Figure S6.

### 3.2. Enzyme-mediated biotransformation of TrOCs

The positive association between DH homolog abundances and treated wastewater percentages suggested that DEET hydrolysis was an enzyme-mediated biotransformation reaction occurring in WW and WW-impacted stream biofilms. We detected the expected DEET hydrolysis transformation product (TP137) in biofilms grown at 80% WW (Figure 1C). The transformation products were consistent with the higher abundance of DH homologs found in these biofilms (Figure 1A). Besides the DEET hydrolysis product, we also detected the product of tolyl group oxidation (i.e., TP222) in 80% WW-grown biofilms, indicating that oxidative biotransformation of DEET also occurred. A previous study on microbial degradation of DEET using activated sludge exclusively reported transformation products of oxidative transformation reactions i.e., amide *N*-deethylation and tolyl group oxidation (Helbling et al., 2010b) (Figure 1B). The *N*-deethylation product, TP164, was not detected in the biofilms probably due to its transient behavior as described previously (Helbling et al., 2010b). In contrast, the amide hydrolysis product DEET detected here (Figure 1B) was only previously reported from a pure culture of *P. putida* isolated from activated sludge through enrichments on 2.6 mM DEET (Rivera-Cancel et al., 2007). To further understand the substrate specificity of this reference *P. putida* DH, we cloned, heterologously expressed, and purified the *P. putida* DH as a reference enzyme, and tested its reactivity against a panel of 183 wastewater-related TrOCs. To avoid cumulative inhibitory effects, the 183 WW-relevant compounds were pooled into 14 sub-mixtures (Table S4 and S5).

Compounds were separated into sub-mixtures using a pooling strategy that aimed to maximize differences in initial preferred biotransformation reactions as predicted by enviPath (Wicker et al., 2016; Zhang and Fenner, 2023). Results showed that the *P. putida* DH transformed 13 out of the 183 TrOCs tested compared to abiotic and inactivated enzyme controls (Figure 1D). With the exceptions of mesalazine, duloxetine, and mefenamic acid, all other compounds biotransformed by the *P. putida* DH contained at least one hydrolyzable moiety (Figure S6). Of those, one compound contained an ester group as sole hydrolyzable moiety, four more compounds contained a combination of esters and one or more amide groups, and five compounds contained exclusively secondary or tertiary amides as hydrolyzable moieties. These results expand on previous findings (Rivera-Cancel et al., 2012) that in addition to the ability of the *P. putida* DH to catalyze hydrolysis of certain tertiary amides, it also exhibits promiscuous activity towards ester and secondary amide-containing contaminants (Figure 1D and Figure S6). Here we use the term ‘promiscuous activity’ to refer to substrate promiscuity, which is the capability of an enzyme to accept other substrates beyond its primary substrate. Of the five compounds for which hydrolysis could exclusively have taken place at the amide functional group (i.e., DEET, MMclB, moclobemide, indomethacin, and tebutam), four (i.e., DEET, MMclB, moclobemide, and indomethacin) contained benzamide functional groups. This points to DH preferring, but not being limited to, the presence of the benzamide moiety in its substrates. Notably, of the over 80 amide-containing compounds tested in our library, five more compounds—enzalutamide, apremilast, tebufenozide, bentazone, and apalutamide—contained benzamide substructures yet did not undergo significant biotransformation. Closer inspection of their molecular structures revealed that all of these compounds contained an ortho-substitution on the benzamide ring with the exception of tebufenozide. This structural characteristic has been previously described to hinder hydrolysis of secondary amides in activated sludge microbial communities due to steric effects (Helbling et al., 2010a).

### 3.3. Construction of a metagenomic library of DEET hydrolase family homologs

Previous studies on the *P. putida* DH revealed it is a di-domain CocE/NonD family enzyme containing both alpha/beta hydrolase and Xaa-Pro dipeptidyl-peptidase protein domains (Rivera-Cancel et al., 2012, 2007). Referred here for simplicity as the ‘DH family’, enzymes in this diverse family were also previously shown to cleave cocaine (Bresler et al., 2000) and α-amino acid esters (Barends et al., 2006). The broad substrate range of DH family enzymes beyond DEET hydrolysis along with our results from the *P. putida* DH (Figure 1D) suggested the potential for DH homologs in WW-grown stream biofilms to transform other TrOCs beside DEET. Therefore, we turned our attention to uncharacterized DH homologs detected in the 75 metagenomes from this study (Table S1). After metagenomic assembly and gene calling (Attrah et al., 2023), we selected a library of 64 full-length DH homologs based on taxonomic and sequence diversity (Table S2, see SI). Taxonomic annotations for the DH sequences were assigned to total 10 different bacterial phyla: *Acidobacteriota*, *Actinobacteriota*, *Bacteriodota*, *Chloroflexota*, *Firmicutes* (*Bacillota*), *Gemmatimonadota*, *Myxococcota*, *Planctomycetota*, *Proteobacteria* (*Pseudomonadota*), and *Verrucomicrobiota*. Surprisingly, a full-length DH sequence with amino acid sequence identity greater than 40% to the *P. putida* DH reference (Rivera-Cancel et al., 2007) could not be recovered from the assemblies, although we did recover DH homologs from *Pseudomonas* and other related genera. Full-length DH homologs recovered from metagenomes with start and stop codons intact had an average of only 25% amino acid sequence identity with the *P. putida* DH (Table S2). The high level of sequence diversity and general conservation of DH homologs across phyla suggested a more conserved role of DH homologs *e.g.*, in primary metabolism, beyond specialized DEET hydrolysis. In order to test this hypothesis experimentally, we next investigated sequence-structure-function relationships of the metagenomic DH family enzymes (Figure 2).

**Figure 2.**
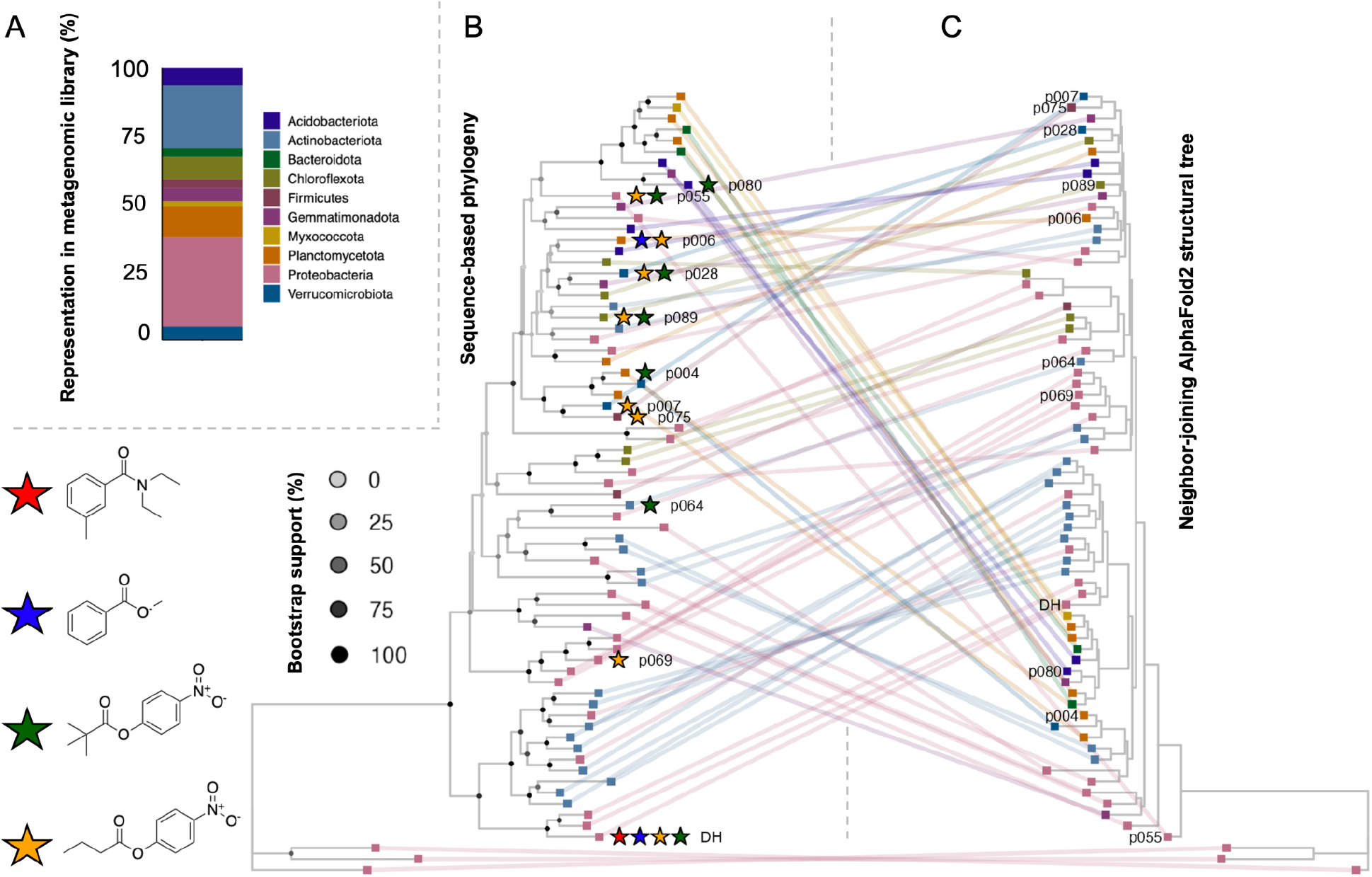
A) Representation of bacterial phyla as a relative abundance within the metagenomic DH homolog library (n = 64) B) Maximum-likelihood sequence-based phylogeny and C) neighbor-joining structural tree of DH family hydrolases. Tips with the same identity are linked between trees. Bootstrap support (1,000 replicates) for internal nodes is indicated by shading. Tips are colored based on the phylum of the assigned taxonomic origin of metagenomic hydrolases. Star colors correspond to activity measured with 4 substrates: Red = DEET (reference compound), Blue = methylbenzoate (aromatic ester), Green = 4-nitrophenyl trimethylacetate (branched aliphatic), Gold = 4-nitrophenyl butyrate (linear aliphatic). Functionally-divergent alpha/beta hydrolases (PDB IDs: 1MJ5, 3HI4 and 6QHV) were used as an outgroup.

### 3.4. Sequence-structure-function relationships among DH family enzymes

We reconstructed a maximum-likelihood phylogeny of metagenomic DH homologs relative to the reference *P. putida* DH (Figure 2A). Although enzymes from many folds are often sequence-diverse, they still often share conserved protein structures (Illergård et al., 2009). Thus, to further test structural similarity of our DH homologs, we used AlphaFold2 (Jumper and Hassabis, 2022) to construct structural models for the 64 metagenomic DH homologs in our library. Structural model comparisons revealed that enzymes shared an average of 1.73 ± 0.43 angstrom (Å) root-mean squared deviation (RMSD) of atomic positions with the *P. putida* reference DH (Table S6). A RMSD less than two indicates relatively high structural similarity and is comparable to values reported in structural alignments of other related members of contaminant-degrading enzyme families (Perez-Garcia et al., 2021; Sidhu et al., 2019). We then compared our sequence phylogeny (Figure 2B) with a neighbor-joining tree generated from the RMSD matrix of all-vs-all structural alignments in our AlphaFold2 library (Figure 2C). This head-to-head tree comparison enabled us to view how clades of sequence-similar DH proteins grouped generally-with structurally-related DH proteins. This approach also enabled us to identify ‘structural deviants’ among the DH homologs discussed further in section 3.6.

Important for this comparison, we verified that our DH homolog structural models were of high confidence with an average predicted local distance difference test (pLDDT) score >80 inclusive of unstructured N- and C-termini (Table S6). Structural alignments of catalytic residues in our library with the *P. putida* DH active site revealed 100% conservation of the catalytic serine and greater than 90% conservation of a complete Ser-His-Asp catalytic triad (Table S6). Our metagenomic library contained more variation (Table S6) with respect to oxyanion hole residues (*P. putida* DH: Y84 and W167) and two other known substrate interacting residues M170 and W214 (Rivera-Cancel et al., 2012). We hypothesized that these active site variants occurring in nature might correspond to altered substrate specificity of the DH homologs in our metagenomic library.

### 3.5. Functional screening of the metagenomic DH homolog library

In order to test the substrate specificity of our metagenomic DH homolog library, we codon optimized, cloned and and expressed each of the 64 metagenomic sequences in *E. coli*. We screened for activity in each of the 64 hydrolase-expressing strains with four different ester or amide-containing substrates (Figure 2): DEET, methylbenzoate, 4-nitrophenyl butyrate, and 4-nitrophenyl trimethylacetate using a combination of HPLC and a previously-optimized 96-well plate protocol (Robinson et al., 2020) (Methods in SI). Eleven out of 64 metagenomic library enzymes tested were identified as active with at least one substrate, primarily with the linear and branched 4-nitrophenyl aliphatic esters (Figure 3). Substrates with ester linkages were preferred by metagenomic DH homologs likely because esters are generally susceptible to nucleophilic attack, e.g., by the catalytic serine of the α/β-fold hydrolase triad conserved among all of the 64 DH homologs.

**Figure 3.**
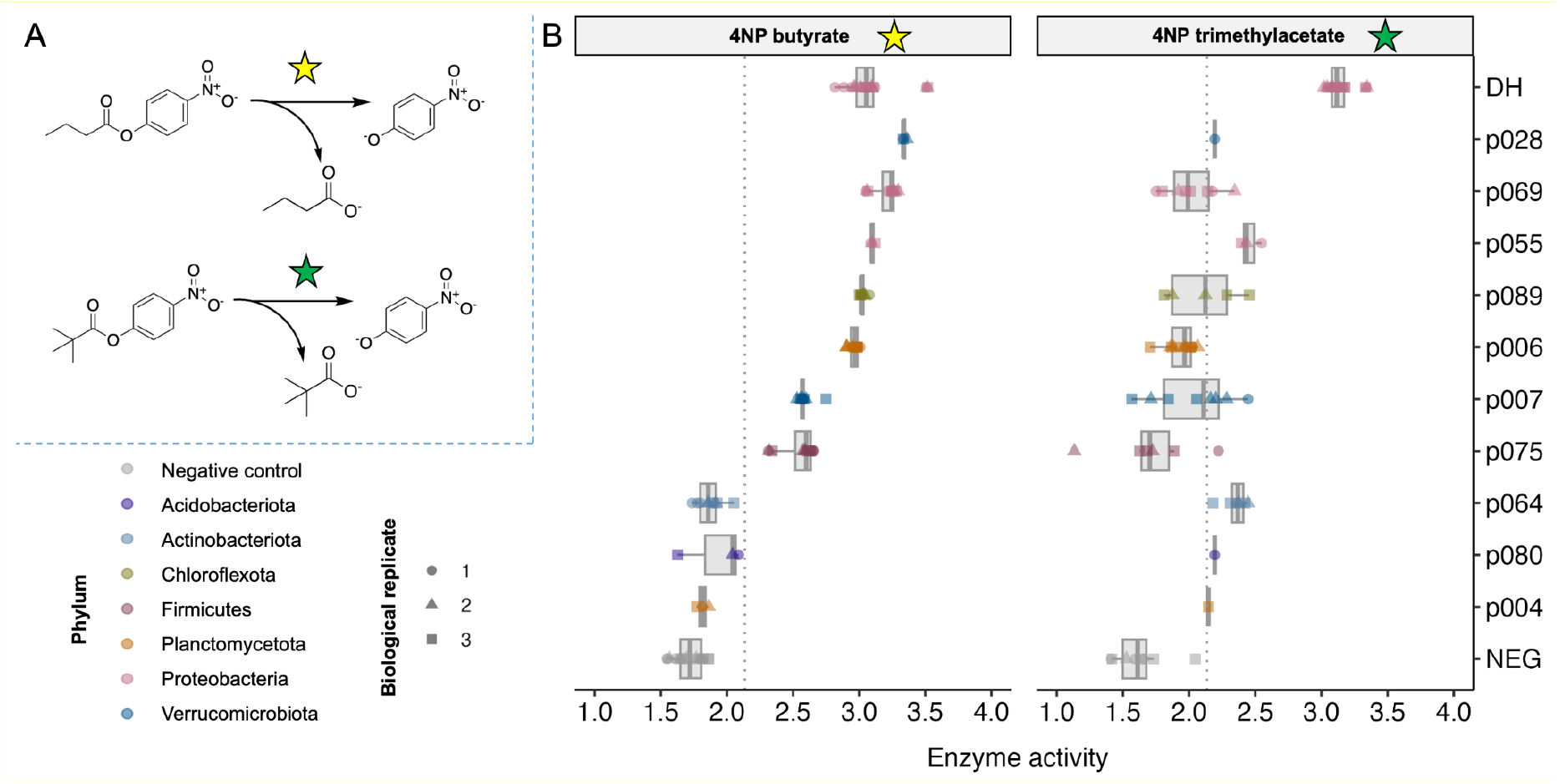
A) Linear and branched 4-nitrophenyl (4NP) ester substrates tested: 4NP butyrate (yellow star) and 4NP trimethylacetate (green star) both release 4NP as a chromophore B) The ten most active metagenomic enzymes in our library tested in a colorimetric screening assay for 4NP hydrolysis activity. Units for enzyme activity are reported as log_10_ of nanomoles of 4NP released per hour by an *E. coli* cell culture with an optical density unit (OD) of 1.0, enabling direct comparison with the activity values reported from previous enzymatic screening studies (Robinson et al., 2020; Smith et al., 2020). Enzyme activity is measured compared to negative control empty vector (NEG) and reference *P. putida* DEET hydrolase (DH). The average activity level calculated across all enzymes and substrates is drawn as a dotted gray line.

Out of the 64 enzymes screened, one metagenomic hydrolase (denoted p006) hydrolyzed methylbenzoate and 4-nitrophenyl butyrate but not DEET (Figure 4). The substrate specificity of p006 is striking in light of previous reports (Rivera-Cancel et al., 2012) confirmed by our own results shown here (Figure 4) that the reference *P. putida* DH hydrolyzes both methylbenzoate and DEET. Compounds containing tertiary amide groups, such as DEET, are more sterically hindered than carboxylic acid esters. Additionally, after testing all 64 DH homologs, surprisingly only the reference *P. putida* DH was capable of DEET hydrolysis (Figure 4). This may be due to the fact that the *P. putida* DH contains active site residues which occur in a unique combination relative to active site residues in the metagenomic DH homologs. Specifically, Y84, W167 and M170 exclusively co-occur in the *P. putida* DH. Swapping each of these residues individually for alanine negatively impacts the kinetics of DEET hydrolysis (Rivera-Cancel et al., 2012). Each of these three residues (Y84, W167 and M170) occurred individually or in pairs in different proteins in our metagenomic DH homolog library. However, all three residues co-occurring together were not identified in any DH homologs (Table S6) in the full dataset or in public databases apart from in the reference *P. putida* DH. This suggests that epistatic interactions between multiple amino acids in the *P. putida* DH active site is critical for DEET hydrolysis. Moreover, in our metagenomic analysis, we identified a naturally-occurring and diverse pool of genes encoding for various combinations of the active site residues Y84, W167 and M170. Such observed genetic diversity within microbial communities indicates a potential for developing specialized functions, especially when exposed to selective pressure from contaminants. Specifically, we propose that different active site variants in this diverse gene pool may contribute to the emergence of specialized degradation enzymes, such as the *P. putida* DH which was identified under DEET enrichment conditions (Rivera-Cancel et al., 2007).

**Figure 4.**
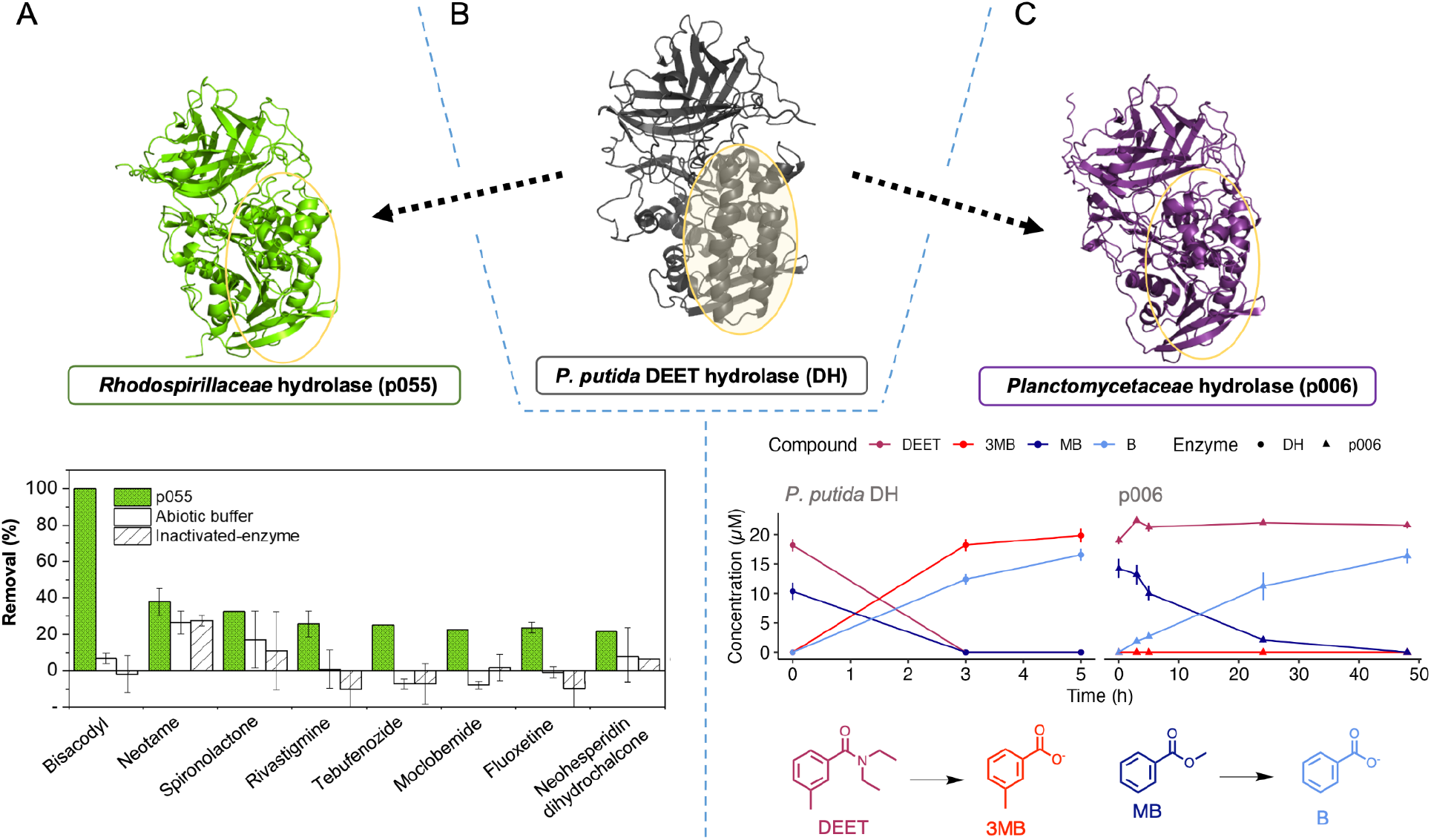
A) AlphaFold2 model of DH homolog p055 and substrate specificity as measured by LC-MS/MS, B) AlphaFold2 model of *P. putida* DH. The yellow encircled region corresponds to an alpha-helix rich region in the vicinity of the active site which has higher alpha helix content than most other active metagenomic DH homologs (Table S7). C) AlphaFold2 model of p006 and HPLC measurements of DEET and methylbenzoate (MB) and product formation of 3-methylbenzoate (red, denoted as “3MB”) and benzoate (light blue, denoted as “B”), respectively, by the *P. putida* DH and p006 (C). Detailed selection criteria and complete biotransformation data are available in Methods and Table S5. Recent analysis of the amino acid sequence p006 suggests a more accurate taxonomic placement to the phylum *’Candidatus Hydrogenedentes*’ based on 93.6% amino acid identity and 100% query cover with proteins from a metagenome-assembled genome (MAG) from wastewater (Lin et al., 2021).

### 3.6. Biochemical characterization of a new metagenomic hydrolase p055 and its reactivity with TrOCs

Among the 64 enzymes in our functional metagenomic library, p055 had the highest average combined enzyme activity with 4-nitrophenyl trimethylacetate and 4-nitrophenyl butyrate. Analysis of the DNA and translated protein sequences of the metagenomic sequence p055 could only resolve the taxonomic origin at the family level as belonging to purple nonsulfur bacteria assigned to the family *Rhodospirillaceae*. By mapping metagenomic raw reads to all full-length DH homologs in the library, we identified p055 as the only active enzyme in our library which also showed a clear increasing trend in metagenomic abundance with WW % (Figure S7).

Furthermore, p055 was ‘structurally deviant’ from the *P. putida* DH and other DH homologs in the metagenomic library (Figure 2). With respect to primary amino acid sequence identity, p055 grouped with other metagenomic hydrolases (Figure 2B), but it was clearly differentiated from the rest of the DH homologs on the basis of its structure (Figure 2C). Closer analysis revealed p055 has much shorter alpha helices in one of its domains relative to reference *P. putida* DH (Figure 4). Of note, the helical regions are in a high-confidence region of the AlphaFold2 models (pLDDT > 95) suggesting the structural deviation was not a modeling artifact. We calculated the overall alpha helix content of the reference *P. putida* DH and found it had higher alpha helix content than all but one other enzyme among the active hydrolases identified in the metagenomic DH homolog library (Table S7). Since the alpha helices in p055 (encircled region, Figure 4) are smaller relative to the *P. putida* DH helices in the vicinity of the active site, we hypothesized p055 structural deviation might also correspond to substrate variation.

To test p055 substrate specificity, we expressed and purified the metagenome-derived enzyme to homogeneity as a 66 kDa protein with a yield of 640 µg/mL (Figure S4). We calculated the specific enzymatic activity in buffers over a range of different pHs and determined p055 had a pH optimum between 8.5 – 9 (Figure S8). The enzyme p055 had an optimal temperature of 35°C (Figure S9) and displayed activity up to 45°C which corresponds well with the optimal growth temperatures of the *Rhodospirillaceae* as the predicted source (Baldani et al., 2014). Following incubation for one hour at 40°C, the enzyme lost 50% of its activity (Figure S10), indicating p055 is not highly thermostable. Loss of activity at 40°C is not unexpected since this enzyme originates from an environmental biofilm grown in WW and stream water where temperatures ranged from 12°C - 14°C (Desiante et al., 2022).

A screening of the substrate promiscuity of this environmental hydrolase p055 with the 183 TrOCs revealed reactivity towards 8 substrates (Figure 5C and Figure S11). By comparing p055 substrate specificity with the *P. putida* DH reference, we identified 20 substrates which were different between the enzymes and two substrates which were shared. Namely, both p055 and *P. putida* DH catalyze the hydrolysis of bisacodyl and moclobemide. Bisacodyl is known as a prodrug that can be metabolized by intestinal and bacterial enzymes to its deacetylated active metabolite. As active esterases, *P. putida* DH and p055 are both capable of facilitating this hydrolytic process.

Overall, we detected activity in approximately 5% of the 626 distinct enzyme-substrate combinations tested. The activity rate with our targeted metagenome mining approach was higher than most untargeted functional metagenomics screens, where on average less than 0.08% of colonies screened are active, depending on the activity screening method (Ferrer et al., 2016). This result highlights the benefits of *in silico* analysis mapping sequence-structure-function relationships to prioritize candidates for metagenomic libraries.

The fact that many of the metagenomic DH homologs tested were active with other organic substrates, yet not active with DEET, also raises the question of whether there are additional responsible enzyme(s) for DEET hydrolysis in stream biofilms. While DH homologs with high similarity to the reference *P. putida* DH were not recovered in the metagenomic assembly process, metagenomic short reads mapped to the *P. putida* DH (Figure S5). Therefore ‘true’ DHs with activity similar to the *P. putida* DH are therefore likely present in low abundance. This also emphasizes the limitations of metagenomic assembly-based approaches for capturing rare, low-abundance genes involved in TrOC biotransformations. Here, we examined the only enzyme family for which we could match an observed TrOC biotransformation to a validated reference sequence detected in our metagenomes. In the environment, we also expect diverse hydrolase families beyond DH and its homologs to contribute to hydrolysis of diverse TrOCs (Hübner et al., 2021), including *downstream effect* compounds. Other DEET biotransformation products e..g, formed from oxidative *N*-deethylation, were observed but the responsible enzymes are currently not well-understood and represent future directions.

It is also worth noting that, within microbial communities, enzymes and their producers exist in a mixture and certain transformation products of a TrOC may be further transformed by enzymes produced by different community members (Wang et al., 2022). All the evidence points to the importance of maintaining (bio)diversity in natural water systems to effectively cope with and challenges by novel entities and anthropogenic activities. In our study, this was exemplified by characterizing new DH homologs spanning ten different microbial phyla from complex WW-impacted biofilm communities. Future studies investigating diverse enzyme families are necessary to expand reference sets for TrOC biodegradation. This will not only deepen our understanding of the fate of TrOCs in environments, but also facilitate the development of more accurate prediction models for assessing TrOC biotransformations in natural and engineered aquatic systems.

## 4. Conclusions

Through combined metagenome mining and experimental characterization, this work yielded new insights into the involvement of DH family enzymes in TrOC biotransformations. Several major conclusions arose from this study:

- This work provided lessons for the computational prediction of responsible enzymes for TrOC removal including the importance of setting a high sequence similarity for assigning protein functions. Depending on enzyme class, functions may or may not be conserved when sequence similarity is low (e.g., less than 40%), even when structural similarity is high as in the case of the DH homologs. Here we characterized enzymes with low amino acid identity to the reference DH. Enzymes bearing low similarity to reference sequences also come from unexplored regions of sequence space thereby presenting opportunities for the discovery of new active metagenomic hydrolases such as p055 with altered substrate specificity for other TrOCs.
- Activity-based screening approaches provide complementary and separate information from sequence- and structure-based approaches to uncover enzymes involved in TrOC biotransformations, as exemplified here. Additional strategies such as machine learning based enzyme-substrate promiscuity predictions or (meta)transcriptomics to identify candidates with differential expression patterns may point towards candidates not detected by shifts in metagenomic abundances.
- Finally, our findings serve as a warning that correlations in ‘omics data do not equal causation. Experimental validation is often critical to test hypotheses generated from microbiome analyses. Metagenomic DH homologs were identified by bioinformatics methods as being active in DEET biotransformation but were not active with DEET, despite being active with other substrates. In addition to sequence similarity, further knowledge on specific amino acids (e.g., active site), secondary structural features (e.g., alpha helices as analyzed here), and experimental or environmental conditions (e.g., pH, temperature) must be incorporated to improve the prediction of TrOC removal in environmental systems.

## CRediT Authorship Contribution Statement

Y.Y.: Conceptualization, Methodology, Investigation, Formal analysis, Writing - Original Draft, Writing - Review & Editing, Visualization; N.F.T.: Methodology, Investigation, Formal analysis, Visualization, Writing - Review & Editing; M.R.S.: Methodology, Formal analysis, Visualization, Writing - Review & Editing; K.F.: Conceptualization, Writing - Review & Editing, Supervision, Funding acquisition; S.L.R.: Conceptualization, Writing - Original Draft, Writing - Review & Editing, Supervision, Funding acquisition.

## Data Availability

Sequencing data are available on NCBI under the BioProject accession number PRJNA1008123. Scripts for data analysis and figures in this manuscript are available on <u>github.com/MSM-group/DEET-hydrolase</u>.

## Acknowledgments

We acknowledge Thomas Fleischmann, René Gall, Max Hofland, and Florian Felder for their technical support. We thank Elia Ceppi and Philipp Longree for assisting with the development of UHPLC-HRMS/MS methods, and Martina Kalt and Miguel Heussi for preparing chemical stock solutions. We thank Kunyang Zhang for assisting with cheminformatics analyses. We thank Giulia Gionchetta, Robert Niederdorfer, and Helmut Bürgmann for their contributions in the metagenomic analysis. We also acknowledge Yasuo Yoshikuni, Ian Blaby, and Miranda Harmon-Smith for their support with metagenomic library constructs. The work (proposal: 10.46936/10.25585/60008420) conducted by the U.S. Department of Energy Joint Genome Institute (https://ror.org/04xm1d337), a DOE Office of Science User Facility, is supported by the Office of Science of the U.S. Department of Energy operated under Contract No. DE-AC02-05CH11231. This work was supported by Eawag Discretionary Funding (for S.L.R. and K.F.) and Swiss National Science Foundation (project no. 200021L_201006, for Y.Y. and K.F.; PZPGP2_209124 for S.L.R.). The graphical abstract was created with a licensed version of BioRender.com.

## Abbreviations

TrOCs: Trace organic contaminants

DEET: *N,N*-diethyl-meta-toluamide

WW: Treated wastewater

DH: DEET hydrolase

TCA: Trichloroacetic acid

DMSO: Dimethylsulfoxide

UHPLC-HRMS/MS: Ultra-High-Performance Liquid Chromatography Coupled to High-Resolution Tandem Mass Spectrometry

FPLC: Fast protein liquid chromatography

