## Supporting Methods and Figures for "Substrate promiscuity of xenobiotic-transforming hydrolases from stream biofilms impacted by treated wastewater"

### Metagenomic sequence prioritization for library construction

The sole characterized reference DEET hydrolase sequence (DH) described in *Pseudomonas putida* (A3R4E0_PSEPU) was used to query 75 metagenomes (Attrah et al. 2023) for homologous sequences using NCBI BLASTp v2.9.0 (Camacho et al., 2009). BLASTp results were filtered to include hits with a bit-score threshold of 50, a query cover of at least 20%, and at least 20% amino acid identity with the query sequence. The resulting 3,500 hits were clustered using CD-HIT (Fu et al., 2012) at 60% amino acid sequence identity cut-off with a word size of 4 to obtain 70 cluster representatives. Cluster representatives were filtered to include only full-length open-reading frames with valid start and stop codons resulting in 64 remaining sequences. To assign the likely taxonomic origin of the 64 DH homologs, the taxonomic classification tool Kaiju was used (Menzel et al., 2016). Since Kaiju did not provide classification for many cases, a reverse BLAST of the metagenomic sequences against the NCBI non-redundant protein database and consensus taxonomic classification based on the identity of the top hit cross-referenced with Kaiju classification (Table S2).

### Short read mapping and coverage calculations

Metagenomic read mapping was performed following the methods described previously (Lee et al., 2021). Mapping of raw metagenomic reads to query DHs was performed using Bowtie2 v2.4.4 (Langmead and Salzberg, 2012), depth was computed using samtools v1.12 (Li et al., 2009), and coverage was calculated using established methods of Albertsen et al. (2013). Bowtie2 parameters used are described in Table S3. Data analysis, statistics and plotting were performed using R within the RStudio integrated development environment (IDE) with the tidyverse library. (R core team, 2022; RStudio Team, 2020; Wickham et al., 2019).

### Phylogenetic analysis

To analyze the phylogenetic relationship of DH family enzymes, 64 DHs in the metagenomic library plus the reference *P. putida* DH and reference cocaine esterase (PDB ID: 3IDA) were aligned using Clustal-Omega v1.2.3 (Sievers and Higgins, 2018). Three outgroup sequences were included as distantly related proteins with the alpha/beta hydrolase fold with PDB IDs: 1MJ5, 3HI4 and 6QHV. A protein phylogenetic tree including 71 sequences was generated from the multiple sequence alignment of DH homologs and reference sequences using IQ-TREE 2 v2.0.3 (Minh et al., 2020). Briefly, IQ-TREE's ModelFinder function was used to identify the top model. ModelFinder identified the best model as LG+F+R5 which is based on an improved general amino acid replacement matrix (Le and Gascuel, 2008). Non-parametric bootstrapping was performed within IQ-TREE with 1,000 replicates.

**Structural modeling and comparison**

Models were assessed for quality by predicted local distance difference test (pLDDT) score and aligned pairwise with the reference *P. putida* structure (AF-A3R4E0-F1) using the PyMOL 'super' command with default parameters. Alpha helix regions of Alphafold structures were extracted using a custom Python script ('helix_analysis.py') and are available on Github (https://github.com/MSM-group/DEET-hydrolase/). To construct a tree of structural relationships, all-vs-all structural comparisons were made and the resulting matrix was converted into a dendrogram using the neighbor-joining algorithm (Saitou and Nei, 1987) implemented in the an 'nj' function in R ape package. Matching nodes between the sequence and structural phylogenetic trees were paired and visualized using the treedataverse library (Yu, 2022).

**Colorimetric activity screening of *P. putida* DH and DH homologs**

Briefly, individual transformants were inoculated in 5 mL lysogeny broth (LB, final concentration 50 mg/L) supplemented with spectinomycin (final conc. 50 µg/mL). Cultures were incubated at 37°C with orbital shaking at 200 rpm until an optimal density of 600 nm (OD600) of 0.5–0.8 was reached. Incubator temperature was then altered to 16°C before adding isopropyl-β-D-thiogalactopyranoside (IPTG, final conc. 0.1 mM) for induction of protein production. The samples were then incubated for 48 hours. Cell pellets were collected by centrifugation of 0.5 mL cell suspensions at 5,000 x g for 10 min and resuspended in 1 mL

Tris-HCl buffer (0.05 M Tris-HCl, pH 8.0). OD600 was then measured to calculate the cell suspension/Tris-HCL buffer ratio in order to normalize the samples to an OD600 of 0.1. Normalized samples were screened for esterase activity using two colorimetric substrates: 4-nitrophenyl butyrate and 4-nitrophenyl trimethylacetate (200 µM final concentration from 8 mM stock solutions in ethanol (EtOH), for a 2% final EtOH in reactions below the limit which would disturb enzyme activity). Individual reactions containing 195 µL normalized cell suspensions and 5 µL substrates were performed in 96-well clear-bottomed plates (Greiner). Standard curves of 4-nitrophenol (8mM stock solutions in EtOH), a positive control of *P. putida* DH, and a negative control of a heat-inactivated DH were included in each plate. Plates were measured using a microplate reader (Synergy H1, BioTek) at 37°C using a continuous measurement program of 60-time point measurements with a time interval of one minute at OD 410 nm.

**Chemicals for metagenomic hydrolases functional screening**

Four substrates for the initial metagenomic hydrolase functional screening were either purchased from Sigma-Aldrich (for 3-methylbenzoic acid, DEET, and sodium benzoate) and Merck (for methylbenzoate). All stock solutions were prepared to 10 mM concentrations in EtOH and proportionally diluted to desired concentrations for activity screening experiments. For the subsequent screening with 183 organic pollutants…

**High-performance liquid chromatography (HPLC)**

Protein expression was conducted as described in the colorimetric activity screening section but with 400 mL culture volumes. Cell pellets were harvested and resuspended in Buffer A (20 mM imidazole, 20 mM Tris-HCl, 500 mM NaCl, 10% (v/v) glycerol). Cell suspensions were then lysed using a sonicator (SONOPULS HD 3200, BANDELIN) with a 3 mm probe (SONOPULS MS73, BANDELIN) at a frequency of 20 kHz. The sample vials were kept in an ice-water bath to prevent sample overheating caused by sonication. Cell lysates were then centrifuged for 30 min at 17,000 x g at 4°C. The ability of cell lysates to biotransform four chemicals (i.e., 3-methylbenzoic acid, sodium benzoate, methylbenzoate, and DEET) was measured on a Dionex Summit system equipped with a Dionex P680 pump, a Dionex ASI-100 autosampler, a UVD 340U photodiode detector, and an UltiMate 3000 column compartment. Briefly, 10 μL samples were loaded onto a Macherey Nagel HPLC column (particle size 5 μm, 150 x 3.0 mm, EC Nuceloshell RP 18 plus) and eluted with 20 mM sodium phosphate monobasic buffer (A) and $H_2O$/acetonitrile (10%/90%) (B) (both amended with 0.04% phosphoric acid, pH ~ 3) at a flow rate of 800 μL/min, with a following gradient: 100% A: 0 –24 min, 100% B: 24 –24.2 min, 100% A 24.2 –28 min. For UV detection, detector wavelengths were set at 210, 220 and 230 nm. Chromeleon (version 6.80) was used for data acquisition and analysis.

**UHPLC-HRMS/MS analysis**

Briefly, for UHPLC analysis, 25 μL samples were loaded onto an ACQUITY Premier BEH C18 Column (particle size 1.7 μm, 100 × 2.1 mm, Waters) and eluted with nano-pure water (A) and methanol (B) (both amended with 0.1% formic acid) at a flow rate of 300 μL/min, with a gradient as follows: 95% A: 0 – 1.5 min, 95% – 5% A:1 – 7.5 min, 5% A: 7.5 – 9.5 min, and 95% A: .5 – 11.5 min. For HRMS, mass spectra were acquired in full scan mode at a resolution of 70,000 at m/z 200 and a scan range of m/z 100 − 1000 in positive/negative switching mode with electrospray ionization (ESI).

**Benchtop protein purification**

For benchtop purification, supernatants were applied to a column packed with Ni-NTA agarose beads (Qiagen) pre-equilibrated in buffer A. The column was subsequently washed with buffer A and then subjected to two washes with buffer B (comprising 40 mM Tris-HCl, 500 mM NaCl, and 10% (v/v) glycerol). Enzymes were eluted from the column using buffer C (containing 500 mM imidazole, 500 mM NaCl, and 10% (v/v) glycerol). Fractions obtained from both benchtop and FPLC purifications were subjected to analysis by SDS-PAGE (Figure S2 – S4). Fractions containing the enzymes of interest, consistent with their expected molecular weights, were eluted by removing imidazole and then underwent buffer exchange into TNG buffer (50 mM Tris-HCl at pH 7.4, 0.1 M NaCl, and 10% (v/v) glycerol). The enzymes were aliquoted, flash-frozen, and stored at –80°C.

**Fast Protein Liquid Chromatography (FPLC) protein purification**

Purification of enzyme candidates were performed using an ÄKTA pure chromatography system (Cytiva) equipped with an S9 sample pump, a U9-M UV monitor and a F9-C fraction collector. Crude extracts were loaded into a HisTrap FF column (5 mL, Ni Sepharose 6 Fast Flow), and measured under 280 nm wavelength. The system were operated at a flow rate of 2.5 mL/min with Buffer A (20 mM Imidazole Tris-HCl, 500 mM NaCl and 10% (v/v) glycerol) and Buffer C (500 mM Imidazole Tris-HCl, 500 mM NaCl and 10% (v/v) glycerol), followed by method listed below. Software Unicorn (version 7.7) was used for data acquisition and analysis.

| Conditions | Volume | Running buffer |
|---|---|---|
| Equilibration | 5 CV* | 100% buffer A |
| Sample application | Total sample volume | |
| Column wash | 20 CV | 100% buffer A |
| Elution buffer | 5 CV | 100% buffer C (step) |
| Equilibration | 5 CV | 100% buffer A |

*CV: column volume

**Supporting Tables**

All supporting tables are uploaded separately in an associated Excel file.

**Supporting Figures**

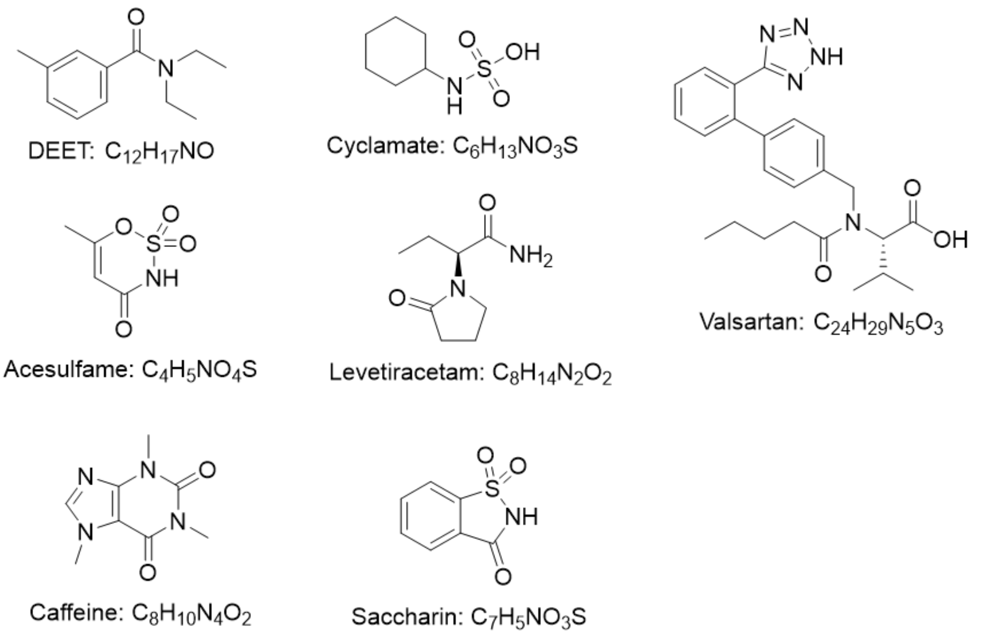

**Figure S1**. Structures of 7 TrOCs (DEET, acesulfame, caffeine, cyclamate, levetiracetam, saccharin and valsartan) that showed the "*downstream effect.*"

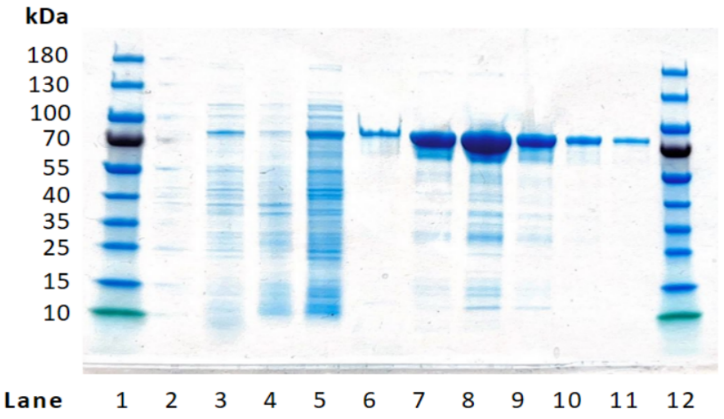


**Figure S2**. SDS-PAGE of purified *P. putida* DEET hydrolase. The protein markers of 180 kDa (Thermo Scientific™ PageRuler™) are shown in lane 1 and 12. Lanes 2 to 11 display various fractions of the ÄKTA pure purification process after expressing the 74 kDa size fusion protein DEET hydrolase (69 kDa w/o tag). The presence of relatively pure DEET hydrolase can be observed in lanes 6-11.

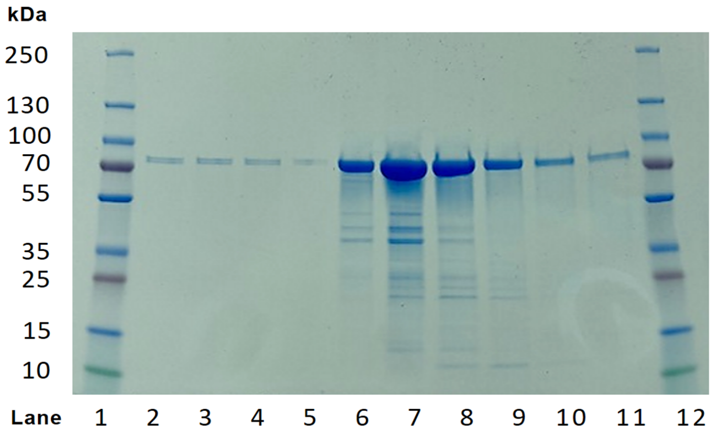


**Figure S3**. SDS-PAGE of purified p006. The protein markers of 250 kDa (Thermo Scientific™ PageRuler™) are shown in lane 1 and 12. Lanes 2 to 11 display various fractions of the ÄKTA pure purification process after expressing the 70 kDa size fusion protein p006 (65 kDa w/o tag). Lanes 6 to 11 show the presence of a pure and relatively high amount of p006.

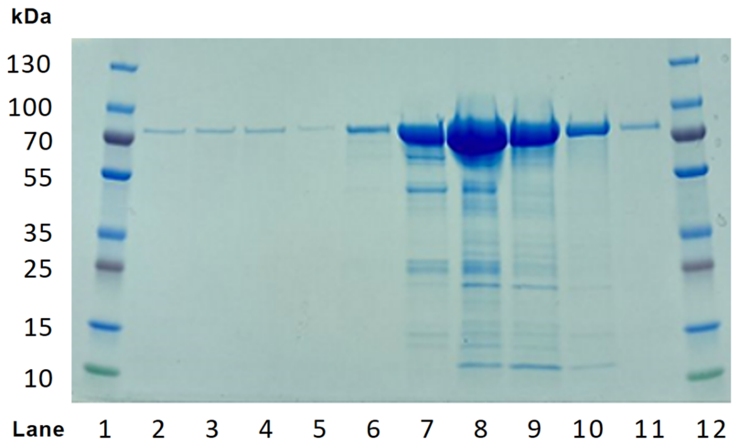


**Figure S4**. SDS-PAGE of purified p055. The protein markers of 250 kDa (Thermo Scientific™ PageRuler™, cropped) are shown in lanes 1 and 12. Lanes 2 to 11 display various fractions of the Äkta pure purification process after expressing the 70 kDa size fusion protein p055 (66 kDa w/o tag). Lanes 6 to 11 show the presence of a pure and relatively high amount of p055.

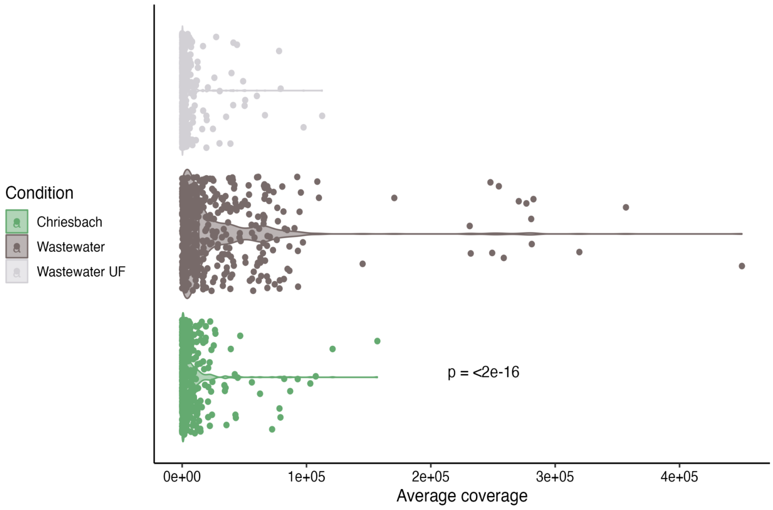


**Figure S5**. Short metagenomic read mapping to the full-length reference *P. putida* DEET hydrolase and coverage calculation. Kruskal-Wallis test yielded a significant difference (*p*-value < 2e-16) in metagenomic read abundances mapping to the *P. putida* DH between treated wastewater (dark gray) vs. stream water (green, peri-urban stream water pumped from the Chriesbach, Dübendorf, CH) and ultrafiltered treated wastewater (light gray).

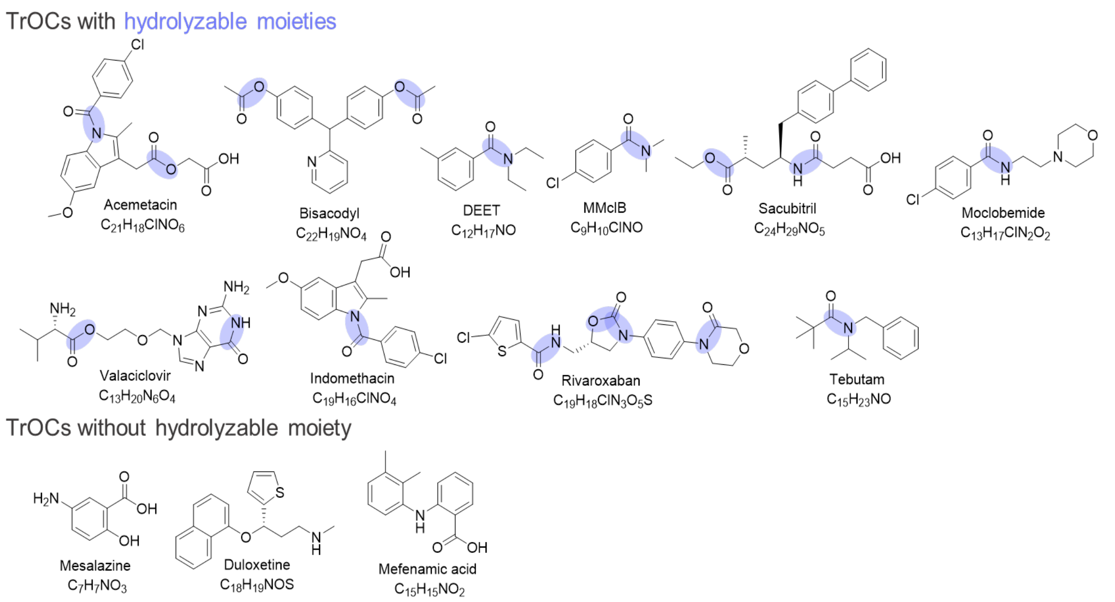


**Figure S6**. Structures of selected substrates transformed by the *P. putida* DH. Detailed selection criteria are included in Methods, complete biotransformation data is available in Table S4.

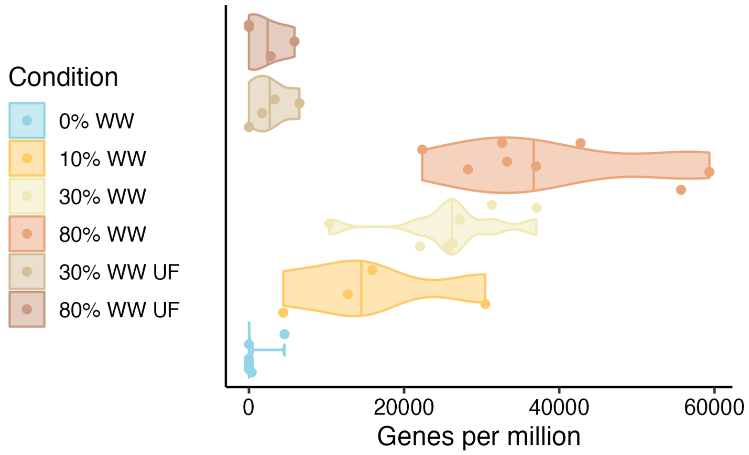


**Figure S7**. Short metagenomic read mapping to the full-length metagenomic hydrolase p055 and coverage calculation across conditions. Kruskal-Wallis test yielded a significant difference between 0% WW and 80% WW (*p*-value < 1.1 e-08).

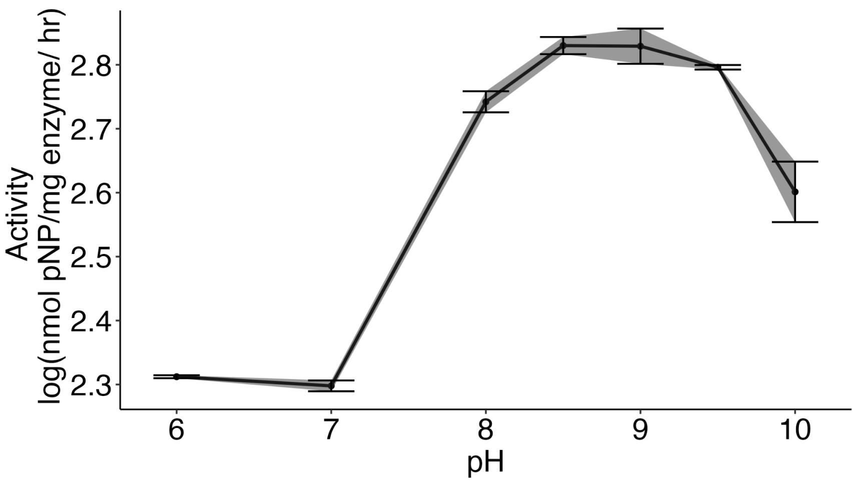


**Figure S8**. Enzymatic activity of p055 reported in terms of nanomoles of *para*-nitrophenol (pNP) released per milligram of purified enzyme per hour over a range of pHs tested (6-10).

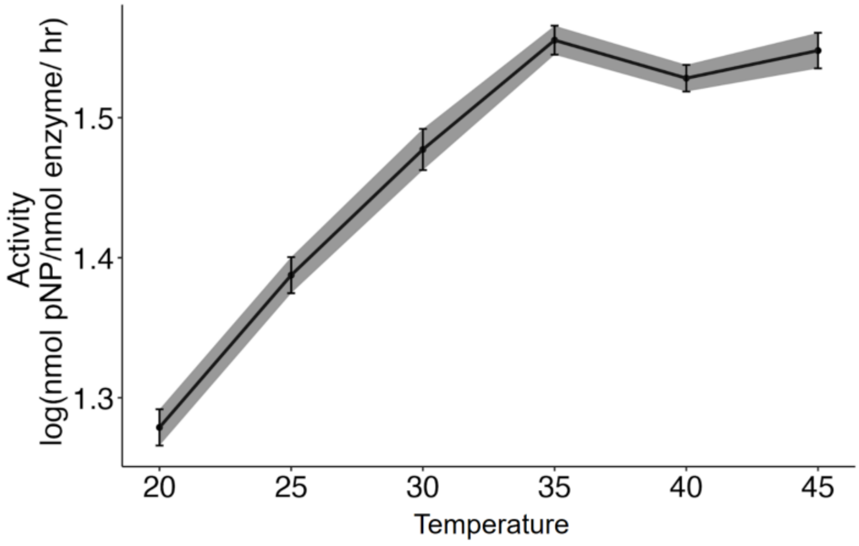


**Figure S9.** Optimal temperature test of enzymatic activity for p055 reported in terms of nanomoles of *para*-nitrophenol (pNP) released per milligram of purified enzyme per hour. The metagenomic DH homolog p055 was tested for hydrolysis activity at different temperatures from 20 - 45°C.

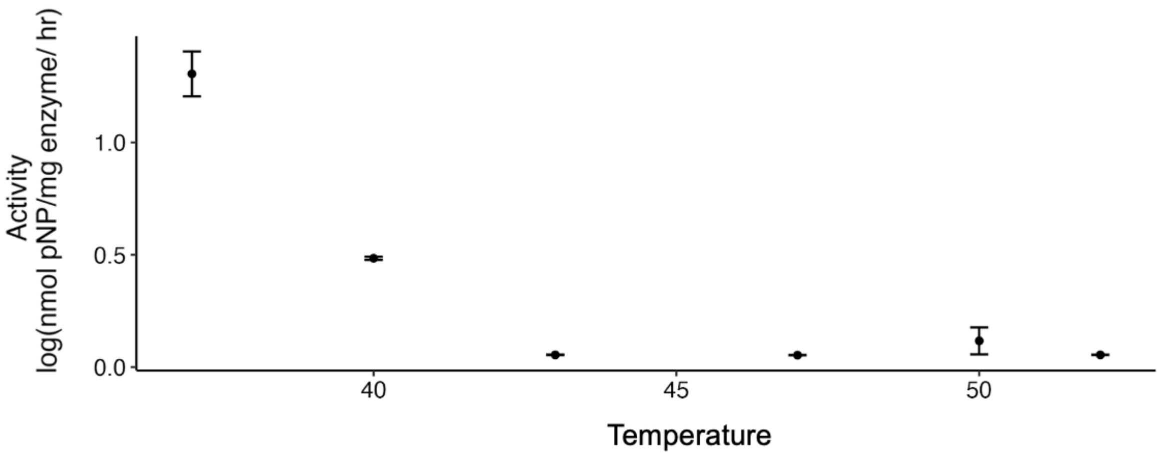


**Figure S10.** Thermostability of p055 reported in terms of nanomoles of *para*-nitrophenol (pNP) released per milligram of purified enzyme per hour. Enzymatic activity at room temperature assessed after incubation for one hour at temperatures ranging from 35°C to 55°C.

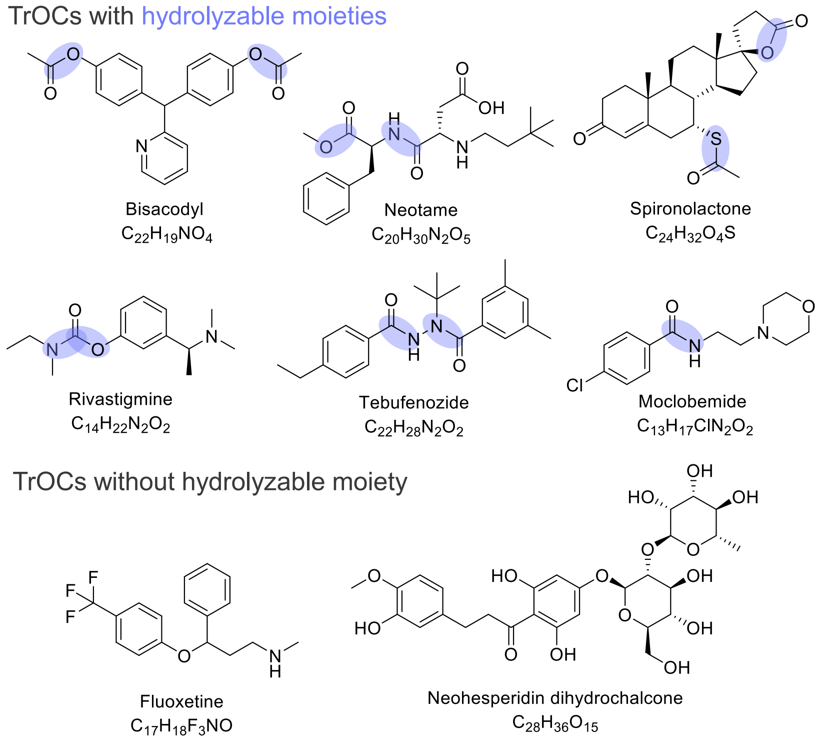


**Figure S11**. Structures of selected substrates transformed by the p055. Detailed selection criteria are included in Methods, complete biotransformation data is available in Table S5.

R core team, 2022. R: A language and environment for statistical [WWW Document]. The R

Project for Statistical Computing. URL https://www.R-project.org/

RStudio Team, 2020. RStudio: Integrated Development Environment for R [WWW Document]. URL http://www.rstudio.com/

Saitou, N., Nei, M., 1987. The neighbor-joining method: a new method for reconstructing phylogenetic trees. Mol. Biol. Evol. 4, 406–425.

Sievers, F., Higgins, D.G., 2018. Clustal Omega for making accurate alignments of many protein sequences. Protein Sci. 27, 135–145.

Wickham, H., Averick, M., Bryan, J., Chang, W., McGowan, L., François, R., Grolemund, G., Hayes, A., Henry, L., Hester, J., Kuhn, M., Pedersen, T., Miller, E., Bache, S., Müller, K., Ooms, J., Robinson, D., Seidel, D., Spinu, V., Takahashi, K., Vaughan, D., Wilke, C., Woo, K., Yutani, H., 2019. Welcome to the tidyverse. J. Open Source Softw. 4, 1686.

Yu, G. (2022). Data Integration, Manipulation and Visualization of Phylogenetic Trees (1st ed.). Chapman and Hall/CRC.
